# Identification and Antibiogram Assay of *Escherichia coli* Isolated from Chicken Eggs

**DOI:** 10.64898/2026.08.08.743651

**Authors:** Rubina Khatun, Md. Rimon Bhuiyan, Mst. Nahida Akter, Nishe Saha, Sharmin afroz, Anna Purnna Ray, K.M. Mozaffor Hossain

## Abstract

**Background:** Escherichia coli contamination of chicken eggs is an important food-safety concern, while antimicrobial-resistant E. coli may contribute to the dissemination of antimicrobial resistance through the food chain. However, information on egg-associated E. coli and its antimicrobial susceptibility in Natore District, Bangladesh, is limited.

**Objectives:** This study aimed to determine the prevalence of E. coli in chicken eggs collected from commercial farms, markets and indigenous/backyard flocks in Natore District, identify the isolates based on cultural, morphological and biochemical characteristics, and assess their antimicrobial susceptibility.

**Materials and Methods:** A total of 84 egg-shell swab samples, comprising 28 samples each from commercial farms, markets and indigenous chicken flocks, were collected from seven upazillas of Natore District between January and June 2023. Samples were cultured on selective and differential media, and presumptive isolates were confirmed by Gram staining, motility and biochemical tests. Antimicrobial susceptibility was determined using the Kirby–Bauer disc-diffusion method against seven antimicrobial agents.

**Results:** E. coli was detected in 56/84 (66.67%) egg samples. Prevalence was highest in indigenous eggs (22/28, 78.57%), followed by farm eggs (18/28, 64.28%) and market eggs (16/28, 57.14%). Among 22 confirmed isolates tested for antimicrobial susceptibility, resistance was highest to neomycin (90.91%) and erythromycin (86.36%), followed by oxytetracycline (77.27%), amoxicillin (68.18%), ciprofloxacin (63.63%), levofloxacin (59.09%) and doxycycline (36.36%). No isolate was sensitive to neomycin or erythromycin.

**Conclusion:** The high prevalence of E. coli and substantial antimicrobial resistance among egg-associated isolates indicate an important food-safety and public-health concern. Improved hygienic egg handling, prudent antimicrobial use and continued antimicrobial-resistance surveillance are warranted throughout the poultry production and marketing chain.

## 1. INTRODUCTION

Eggs are a delicious, low-calorie and highly nutritious food that provides high-biological-value protein and a well-balanced range of nutrients suitable for people of all ages (Watkins, 2017). The poultry sector in Bangladesh produces two broad categories of chickens, one for eggs and one for meat, and in FY 2023-2024 total national egg production reached 2,374.97 crore, about 105.04% of the predicted target, according to the Department of Livestock Services. The avian egg is a valuable source of protein, lipids, vitamins, minerals and growth factors required by the developing embryo, and various biological activities of eggs and egg components-including novel antimicrobial, anti-adhesive, immune-modulatory, anticancer, antihypertensive and antioxidant properties-have drawn attention to their significance for human health and disease prevention (Kovacs-Nolan et al., 2005). However, eggs contaminated with dangerous bacteria can also be harmful to health.

Food poisoning caused by egg-borne pathogens can cause vomiting, nausea, diarrhoea and abdominal cramps, in addition to serious morbidity or death, because the nutrient-rich environment inside eggs is ideal for bacterial growth. Numerous bacteria, including *Salmonella*, *Escherichia coli*, *Proteus*, *Listeria monocytogenes*, *Staphylococcus*, *Streptococcus* and *Bacillus*, can contaminate eggs (Lee et al., 2016). *E. coli* is a common member of the gut microbiota of farm animals, poultry and humans; while most isolates are harmless, certain strains are pathogenic and can cause severe illness in people who consume contaminated food (Begum et al., 2014). Over the past two decades numerous outbreaks of gastrointestinal illness caused by food-borne pathogenic *E. coli*, especially O157:H7, have been reported (Armstrong et al., 1996). Although mechanical processes can transfer bacteria into eggs, *E. coli* typically contaminates the egg surface (Safaei et al., 2011); cracked eggs, soiled shells and storage in contaminated environments increase the likelihood of contamination, and once the shell breaks the contents are more likely to be affected (Neira et al., 2017). Most eggs are sterile when laid, and bacterial contamination generally occurs afterwards, when the egg contacts dirt, faeces, dust, trays or nest material (Board and Tranter, 1995).

*E. coli* is also a common cause of colibacillosis in commercial poultry, manifesting as septicaemia, pericarditis, airsacculitis and mortality-accounting for about 28% of deaths in Sonali flocks (Biswas et al., 2006). It is a Gram-negative, aerobic, rod-shaped, flagellated, motile, oxidase-negative, non-spore-forming member of the family Enterobacteriaceae capable of producing endotoxins (Buxton and Fraser, 1977). Pathogenic *E. coli* is broadly grouped into enteropathogenic and uropathogenic types, with further classification into enterotoxigenic (ETEC), enteropathogenic (EPEC), enteroinvasive (EIEC), enteroaggregative (EAggEC) and enterohaemorrhagic (EHEC) pathotypes (Nataro and Kaper, 1998).

Antibiotic therapy is considered the principal driver of the emergence, selection and spread of antimicrobial-resistant organisms in both human and veterinary medicine (Witte, 1998; Neu, 1992), even though antibiotics remain the main tool for reducing the incidence and mortality of avian colibacillosis (Watts et al., 1993). In Bangladesh, antimicrobials such as ciprofloxacin, streptomycin, gentamicin, erythromycin, tetracycline and, occasionally, furazolidone are authorised and widely used to treat *E. coli* infections in broilers and layers. The consequent development of antimicrobial resistance is a major global public-health concern (Kaye et al., 2004); resistant bacteria typically acquire resistance through chromosomal mutation and the acquisition of multidrug-resistant plasmids (Finch et al., 2003). A cross-sectional survey in Bangladesh found that 98% of commercial chicken farms used antimicrobials in their current production cycle and 85% of farmers administered them prophylactically (Imam et al., 2020). Antimicrobial exercise and abuse are considered the principal drivers selecting for resistance in bacteria (Okeke et al., 1999; Moreno et al., 2000), and although antibiotic resistance is a natural, ultimately unavoidable phenomenon, most of its recent increase is attributable to overuse and misuse of antimicrobials (McGowan, 1983; Arnold and Straus, 2005). Careful, judicious use of antibiotics can prolong the useful life of existing and future medicines and may delay the emergence of clinically important resistance (Dowell et al., 1998).

The continuous selection pressure exerted by widely used antibiotics is largely responsible for the rise of multi-resistant bacterial strains (Kolar et al., 2001), a problem compounded by inappropriate prescribing and misuse of these drugs (Imam et al., 2020). When an antibiotic is used against a bacterial infection, susceptible organisms-including components of the normal microbiota-are eliminated together with the pathogen, allowing any resistant bacteria present to multiply and become dominant. Bacteria evade antibiotics chiefly through enzymatic inactivation, modification of target binding sites, efflux activity and reduced membrane permeability, and resistance may be intrinsic (chromosomally encoded) or acquired through horizontal gene transfer from other bacteria or the environment (Agyare et al., 2018).

The poultry industry uses antibiotics at sub-therapeutic or therapeutic levels to enhance growth, feed-conversion efficiency and disease control (Arathy et al., 2011). Antibiotic resistance together with the presence of invasion (INVE) and virulence-associated Shiga-like toxin (SLT1/2) genes among emerging pathogenic serotypes of *E. coli* isolated from table eggs is a public-health concern that highlights the need for improved management across production and marketing channels to ensure consumers receive eggs free of *E. coli* (Vinayananda et al., 2017). A comprehensive European resistance-monitoring survey of 1,462 *E. coli* isolates from primary producers, slaughterhouses and retail outlets found resistance to third-generation cephalosporins and fluoroquinolones in addition to well-known classes such as tetracyclines and sulphonamides; 85-93% of isolates from laying hens, chicken meat and turkey meat were resistant to at least one antimicrobial class, and 73-84% to multiple classes, whereas most isolates from broilers were sensitive to all agents tested (Kaesbohrer et al., 2012). *E. coli*, together with *Enterococcus faecalis*, is used worldwide as an indicator organism for monitoring antimicrobial resistance because of its wide distribution in the animal gastrointestinal tract (Wray and Gnanou, 2000).

Among the beta-lactamases produced by Gram-negative bacteria, carbapenemases, AmpC beta-lactamases and extended-spectrum beta-lactamases (ESBLs) are of greatest clinical importance in *E. coli*. ESBLs, mostly plasmid-encoded members of Ambler class A, confer resistance to monobactams, penicillins and first-, second- and third-generation cephalosporins, but not to carbapenems or cephamycins, and are inhibited by clavulanic acid, tazobactam and sulbactam (Philippon et al., 1989). In Bangladesh, *E. coli* has previously been isolated from plants, milk, eggs, water, chicken carcasses and diarrhoeic calves (Hasina, 2006), and as the most prevalent commensal gut bacterium of humans and animals it remains an important zoonotic agent capable of causing infectious disease in both (Costa et al., 2008).

Given the food-safety and public-health importance of egg-borne *E. coli* and the growing threat of antimicrobial resistance, the present study was carried out with the following objectives: (i) to determine the prevalence of *E. coli* isolated from chicken eggs of Natore District, Bangladesh; (ii) to isolate *E. coli* from chicken eggs; (iii) to identify the isolated *E. coli*; (iv) to determine the antibiogram of the isolated *E. coli*; and (v) to evaluate the public-health significance of *E. coli* recovered from chicken eggs of Natore District, Bangladesh.

## 2. MATERIALS AND METHODS

### 2.1 Study area and duration

The study was carried out in the Microbiology Laboratory, Department of Veterinary and Animal Sciences, University of Rajshahi, from January to June 2023. Samples were collected from randomly selected commercial layer farms, sale outlets (wholesalers and retailers) and indigenous chicken flocks in different upazillas of Natore District, Bangladesh.

### 2.2 Sample collection

A total of 84 egg samples were collected from seven upazillas of Natore District (Bagatipara, Baraigram, Gurudaspur, Lalpur, Naldanga, Natore Sadar and Singra) to isolate and identify *E. coli*. Twelve eggs were collected from each upazilla: four from a commercial farm, four from the market (wholesaler or retailer) and four indigenous chicken eggs from village households, giving 28 samples in each of the three source categories. Samples were transported to the Microbiology Laboratory for bacteriological analysis. A sterile swab moistened with sterile saline was rubbed over each egg’s shell surface, and each swab was then immersed in 6 ml of sterile saline as a “shell wash” (Adesiyun et al., 2005); the shell wash was homogenised by mini-vortex and inoculated onto different culture media.

### 2.3 Media, reagents and equipment

Liquid media used included Nutrient Broth, Peptone Water Broth, Methyl Red-Voges-Proskauer (MR-VP) Broth and sugar (carbohydrate) media, while solid media included Nutrient Agar, Blood Agar, MacConkey Agar, Eosin Methylene Blue (EMB) Agar, Triple Sugar Iron (TSI) Agar, Salmonella-Shigella (SS) Agar, Brilliant Green Agar, Mueller-Hinton Agar and Simmons’ Citrate Agar, all prepared according to the manufacturers’ instructions and sterilised by autoclaving (121°C, 15 psi, 15 minutes). Reagents included Gram’s stain (crystal violet, Gram’s iodine, acetone-alcohol, safranin), methyl red solution, Voges-Proskauer (Barritt’s) reagents, Kovac’s reagent, phosphate-buffered saline, physiological saline, phenol red indicator, 50% glycerine and 3% hydrogen peroxide. Standard laboratory glassware and equipment-including a compound microscope, autoclave, hot-air oven, incubator, refrigerator, Durham’s tubes, inoculating loops, Bunsen burner and related consumables-were used throughout.

### 2.4 Isolation of E. coli

Following enrichment in Nutrient Broth, a loopful of culture was streaked onto EMB agar and incubated aerobically at 37°C for 18-24 hours. Colonies with a dark centre and characteristic metallic green sheen were presumptively identified as *E. coli*. Colony morphology was examined and pure cultures were obtained by sub-culturing a well-isolated colony onto fresh EMB agar and incubating at 37°C for 24 hours. A single representative colony from each positive sample was then subjected to a battery of confirmatory tests.

### 2.5 Identification of E. coli

Colony morphology (form, size, surface texture, elevation, margin, colour and opacity) was recorded after 24 hours of incubation. Gram’s staining was performed following standard procedure (Cheesbrough, 1985): a bacterial smear was heat-fixed, stained with crystal violet, mordanted with Gram’s iodine, decolourised with acetone-alcohol, counterstained with safranin and examined under oil immersion at 1000x magnification. Motility was assessed by the hanging-drop technique using a compound microscope. Biochemical identification comprised the indole test, methyl red (MR) test, Voges-Proskauer (VP) test, catalase test, sugar-fermentation reactions with five basic sugars (dextrose, lactose, sucrose, maltose and mannitol) and the Triple Sugar Iron (TSI) agar slant reaction, all performed according to standard methods (Cowan, 1985; Cheesbrough, 1985).

### 2.6 Antibiogram assay

Antimicrobial susceptibility of the confirmed *E. coli* isolates was determined by the Kirby-Bauer disc-diffusion method (Bauer and Shiloach, 1975) on Mueller-Hinton agar, following the guidelines of the Clinical and Laboratory Standards Institute (CLSI, 2018; document M100-S17). Seven commercially available antibiotic discs (Hi-Media, India)-ciprofloxacin (5 µg), amoxicillin (30 µg), doxycycline (30 µg), oxytetracycline (30 µg), levofloxacin (5 µg), neomycin (30 µg) and erythromycin (15 µg)-representing antimicrobial classes commonly used in field veterinary practice were tested. Broth cultures of individual isolates, standardised by overnight incubation at 37°C, were spread evenly over Mueller-Hinton agar plates with a sterile glass spreader, and antibiotic discs were placed on the inoculated surface, each about 1 cm apart, with three to four discs per plate. Plates were incubated at 37°C for 16-18 hours, after which the diameter of the zone of inhibition around each disc was measured to the nearest millimetre with a ruler and interpreted as resistant, intermediate or sensitive according to CLSI (2018) zone-diameter breakpoints (Table 1).

**Table 1.**
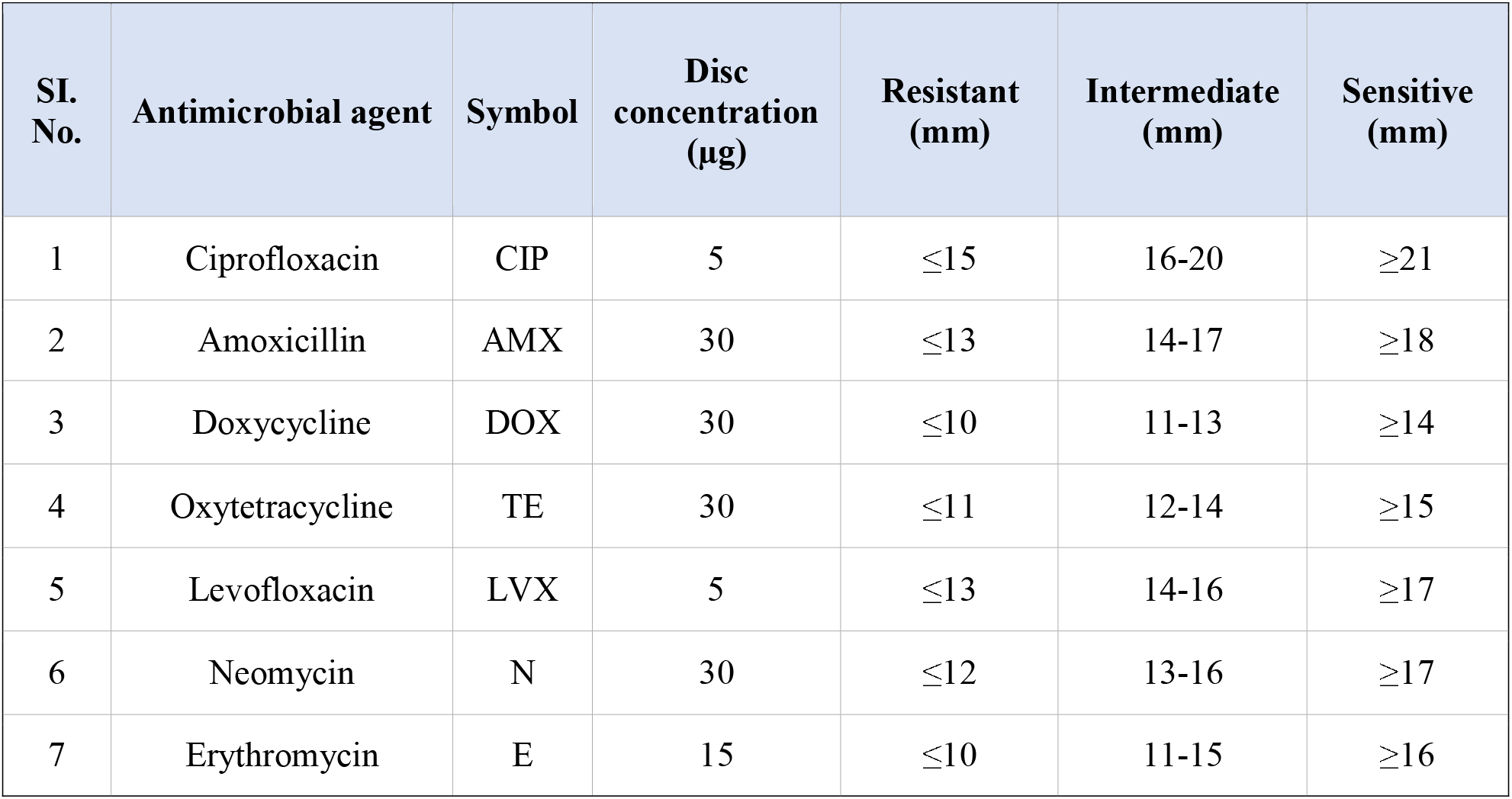
Zone-diameter interpretative standards used for antimicrobial susceptibility testing (CLSI, 2018)

| SI. No. | Antimicrobial agent | Symbol | Disc concentration (µg) | Resistant (mm) | Intermediate (mm) | Sensitive (mm) |
| --- | --- | --- | --- | --- | --- | --- |
| 1 | Ciprofloxacin | CIP | 5 | ≤15 | 16-20 | ≥21 |
| 2 | Amoxicillin | AMX | 30 | ≤13 | 14-17 | ≥18 |
| 3 | Doxycycline | DOX | 30 | ≤10 | 11-13 | ≥14 |
| 4 | Oxytetracycline | TE | 30 | ≤11 | 12-14 | ≥15 |
| 5 | Levofloxacin | LVX | 5 | ≤13 | 14-16 | ≥17 |
| 6 | Neomycin | N | 30 | ≤12 | 13-16 | ≥17 |
| 7 | Erythromycin | E | 15 | ≤10 | 11-15 | ≥16 |

### 2.7 Maintenance of stock cultures

Confirmed isolates were preserved in 50% sterile buffered glycerine (equal parts pure glycerine and phosphate-buffered saline) at -20°C, a method suitable for long-term storage of bacterial isolates without altering their original characteristics.

### 2.8 Data analysis

Data on the prevalence of *E. coli* by source and on antimicrobial susceptibility were compiled and expressed as simple percentages and summarised in tabular form.

## 3. RESULTS

### 3.1 Cultural characteristics

Growth of *E. coli* in Nutrient Broth was indicated by diffuse turbidity and, occasionally, pellicle formation. On Nutrient Agar, smooth, round, white-to-greyish-white colonies developed. On EMB agar, characteristic colonies with a dark centre and a metallic green sheen were produced, and on MacConkey agar the isolates produced bright pink colonies as a result of lactose fermentation. On Brilliant Green Agar, yellow colonies formed with a colour change of the medium from red to yellow, while on XLD agar yellow colonies developed with the medium changing from yellowish-red to yellow, together with partial inhibition of xylose, lactose and sucrose degradation. A summary of the colonial characteristics of the isolates on the different culture media is presented in Table 2.

**Fig. 1.**
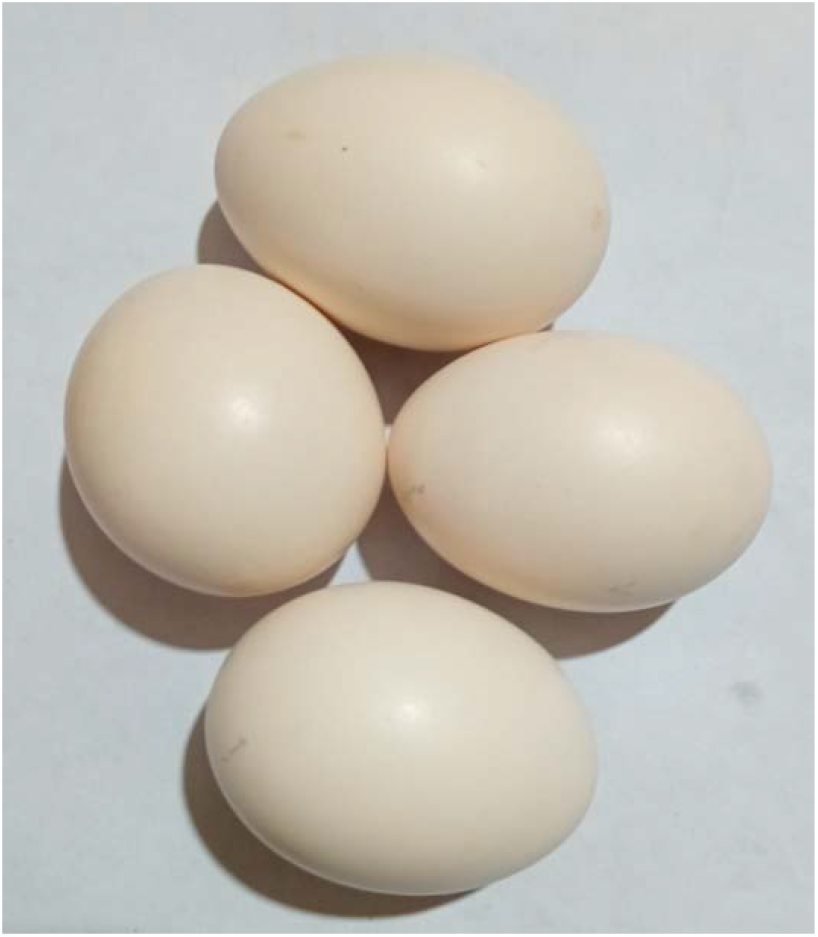
**a.** Farm chicken eggs collected for the study

**Fig. 1.**
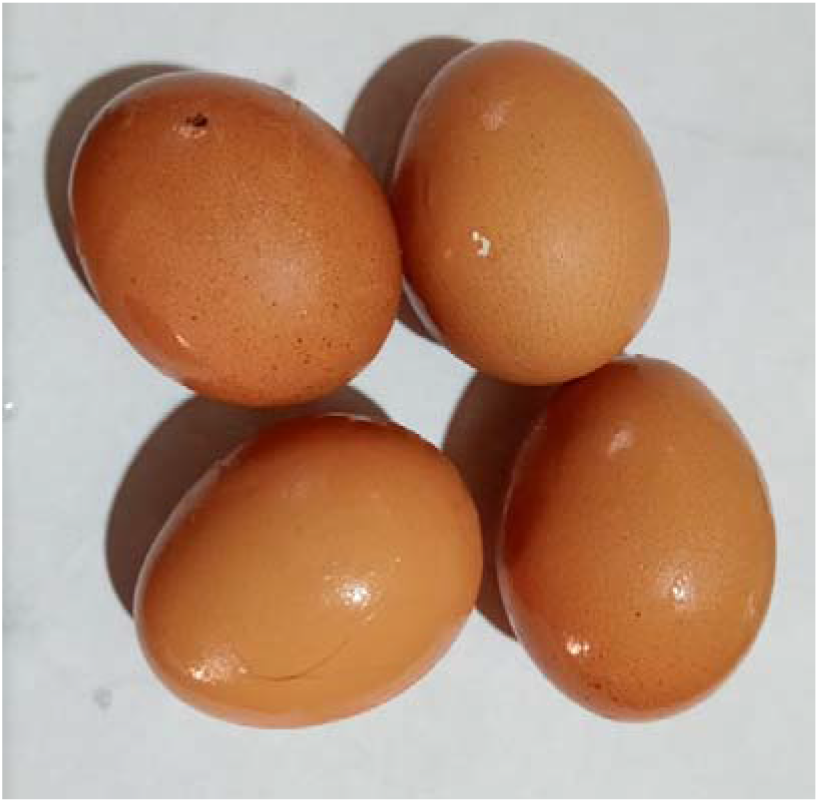
**b.** Indigenous (backyard) chicken eggs collected for the study

**Fig. 2.**
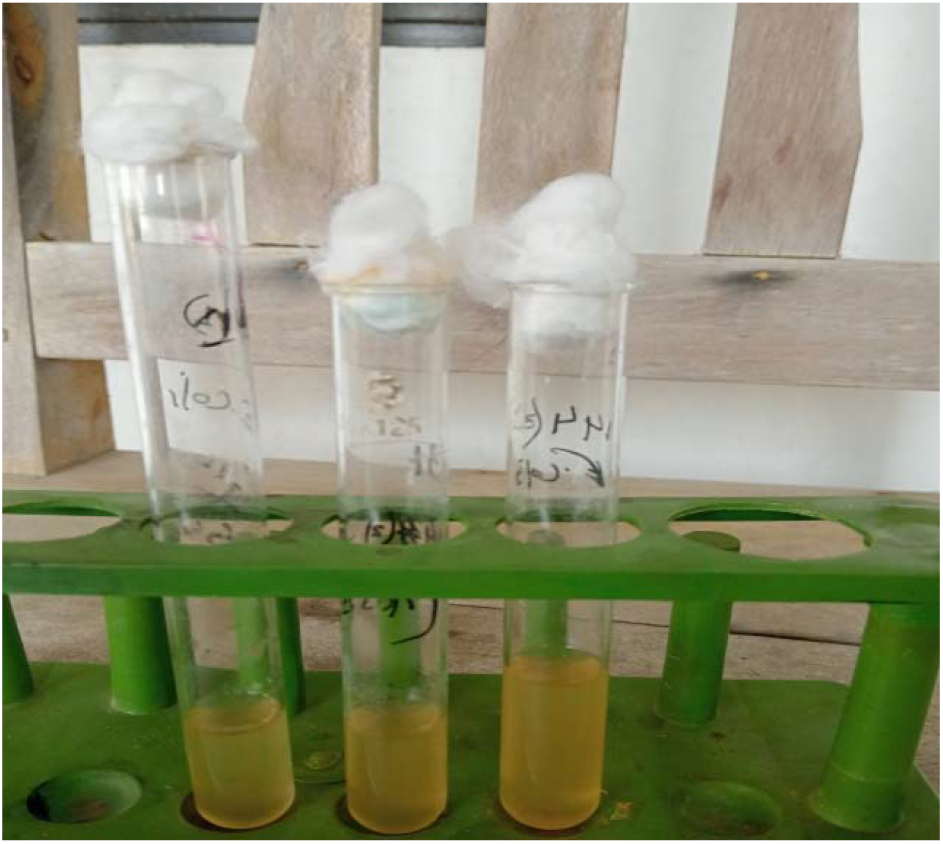
Growth of E. coli in Nutrient Broth showing diffuse turbidity

**Fig. 3.**
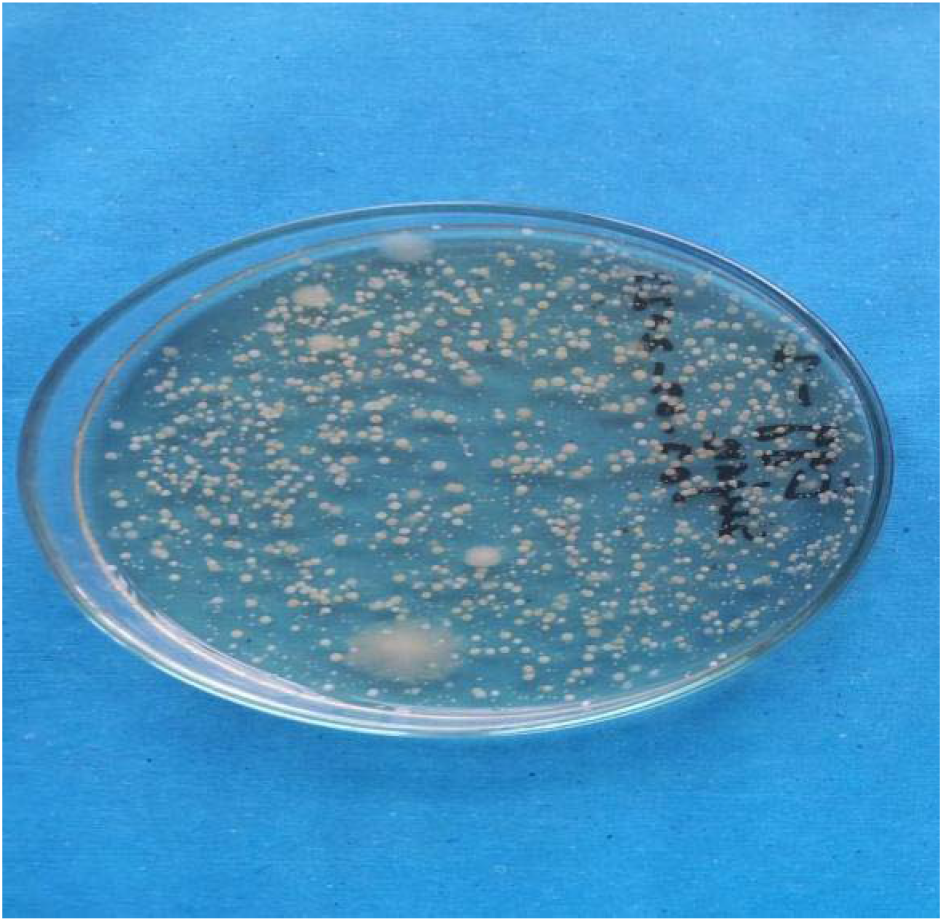
Smooth, white to greyish-white colonies of E. coli on Nutrient Agar

**Fig. 4.**
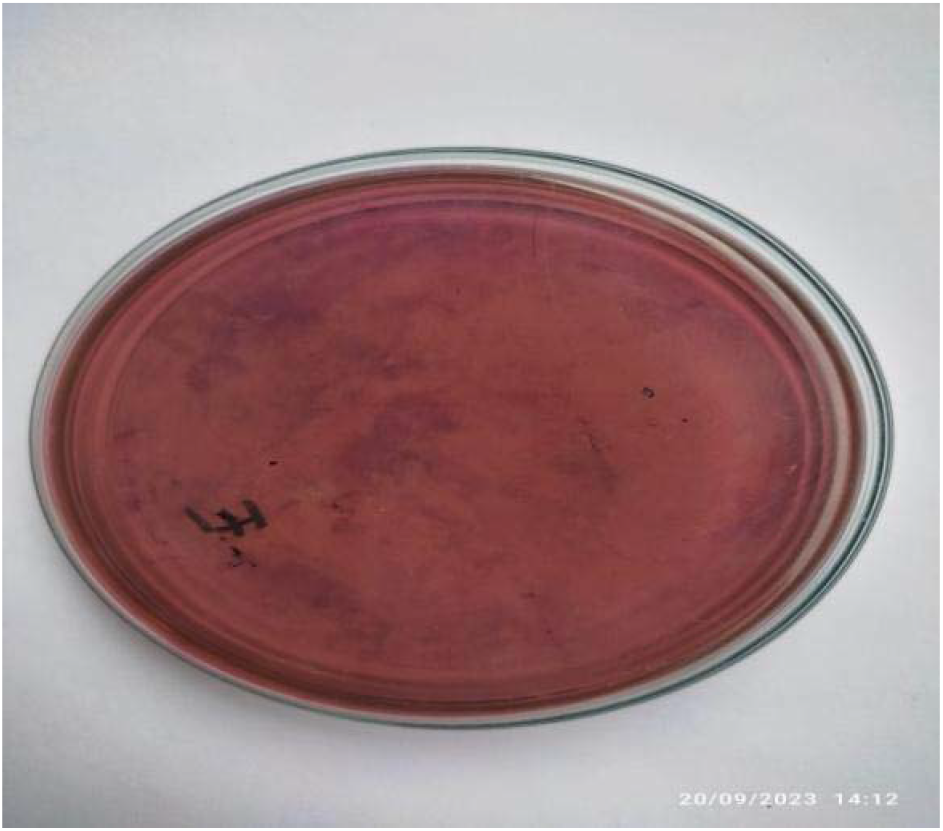
Colonies of E. coli on Eosin Methylene Blue (EMB) Agar

**Fig. 5.**
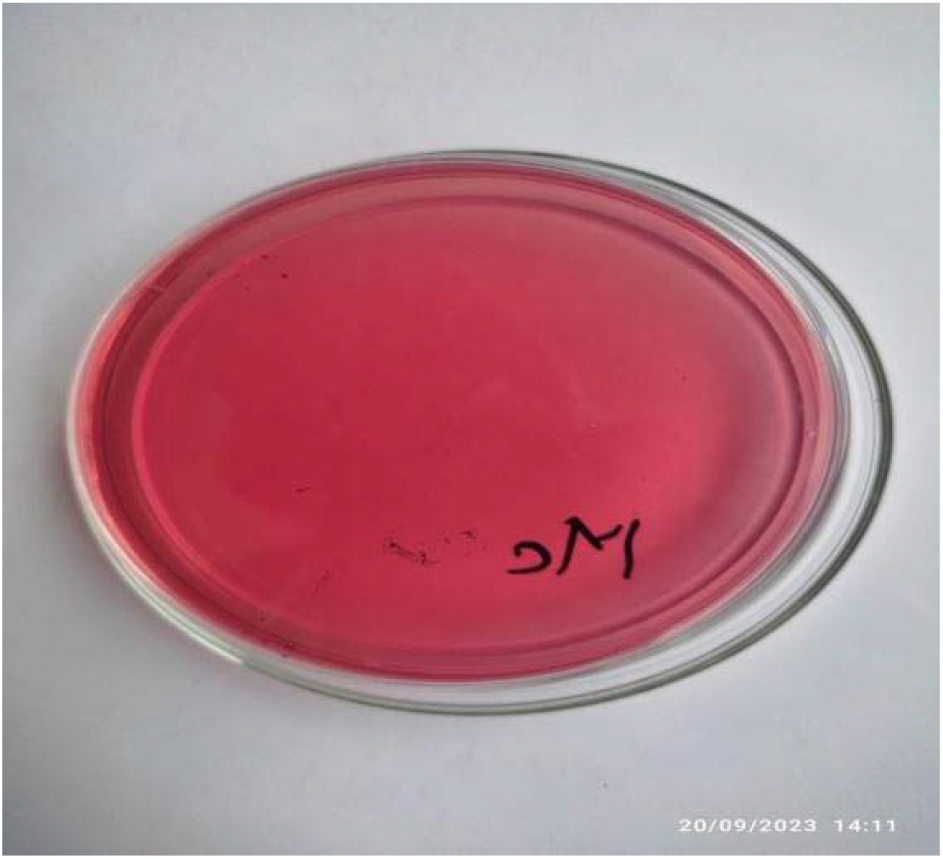
Bright pink colonies of E. coli on MacConkey Agar

**Fig. 6.**
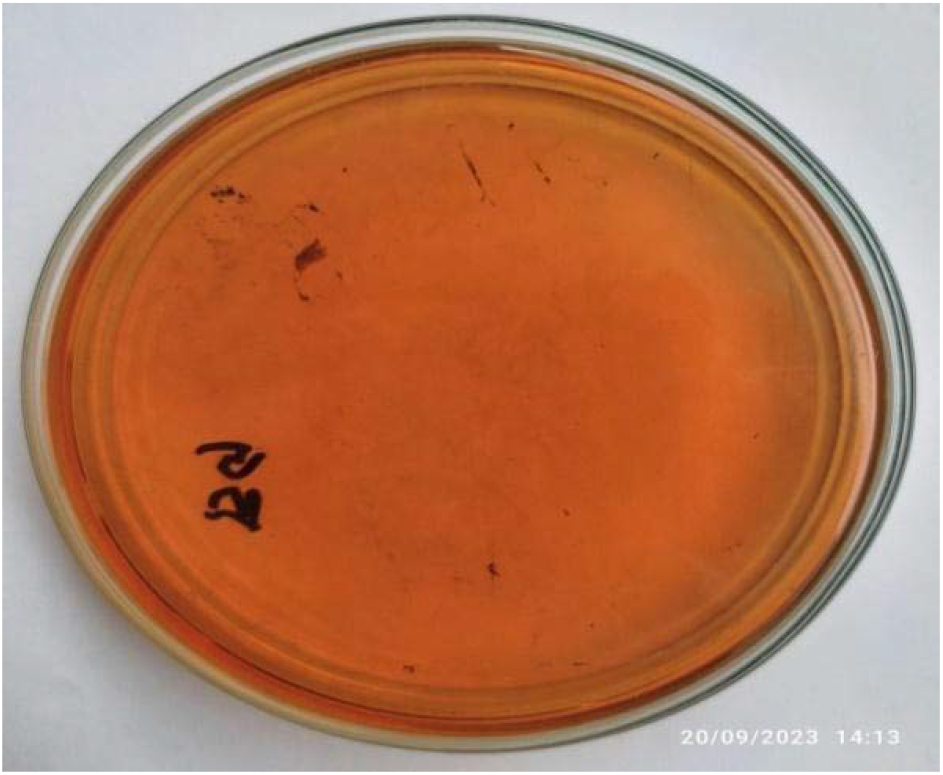
Colonies of E. coli on Brilliant Green Agar

**Table 2.** Cultural, staining and motility characteristics of E. coli isolated from chicken eggs.

| Medium | Colonial characteristics |
| --- | --- |
| Nutrient Agar | Smooth, circular, white to greyish-white colonies |
| EMB Agar | Smooth, circular colonies with a characteristic metallic green sheen |
| MacConkey Agar | Smooth, bright pink colonies (lactose fermenters) |
| Brilliant Green Agar | Yellow colonies; medium changes from red to yellow |
| SS Agar | Slight growth; pink to rose-red colonies |
| XLD Agar | Yellow colonies; medium changes from yellowish-red to yellow |
| Gram's staining | Gram-negative, short plump rods, single or paired |
| Motility test | Motile (positive swinging movement under hanging-drop examination) |

### 3.2 Staining and motility

On microscopic examination of Gram-stained smears, the isolates appeared as small, Gram-negative, rod-shaped organisms occurring singly or in pairs. In the hanging-drop motility test, all isolates displayed active swinging (twitching/darting) movement, confirming that the recovered *E. coli* were motile.

**Fig. 7.**
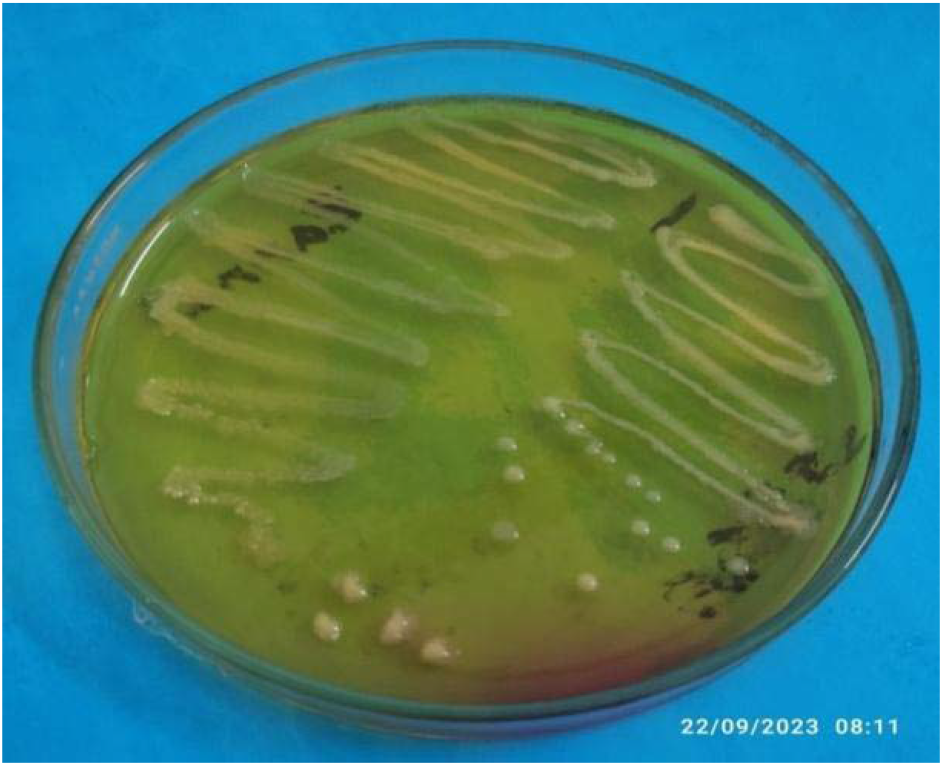
Yellow colonies of E. coli on XLD Agar

### 3.3 Biochemical characterisation

All isolates fermented the five basic sugars tested (dextrose, lactose, sucrose, maltose and mannitol) with the production of both acid and gas, evidenced by a colour change from red to yellow and gas accumulation in the inverted Durham’s tubes. All isolates were indole-positive, catalase-positive and methyl-red-positive, but Voges-Proskauer-negative and Simmons’ citrate-negative. On TSI agar, isolates produced an acidic slant and acidic butt (yellow slant/yellow butt) with gas production and no blackening, indicating fermentation of glucose, lactose and sucrose without hydrogen sulphide production. These biochemical reactions were consistent with the identification of the isolates as *E. coli* (Table 3).

**Table 3.** Biochemical properties of the isolated E. coli.

| Biochemical test | Result |
| --- | --- |
| Sugar fermentation (dextrose, lactose, sucrose, maltose, mannitol) | Positive (acid and gas) for all five sugars |
| Indole test | Positive |
| Catalase test | Positive |
| Methyl red (MR) test | Positive |
| Voges-Proskauer (VP) test | Negative |
| TSI agar slant reaction | Acidic slant and acidic butt with gas, no H<sub>2</sub>S |
| Simmons' citrate utilisation test | Negative |

**Fig. 8.**
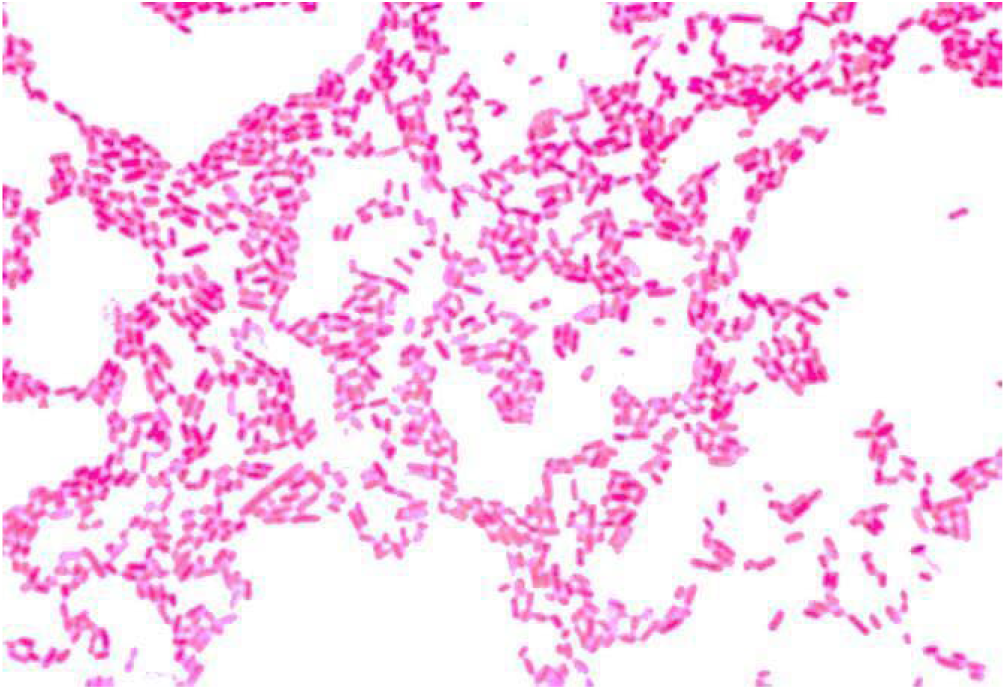
Gram’s-stained smear showing Gram-negative, rod-shaped E. coli (single/paired arrangement)

**Fig. 9.**
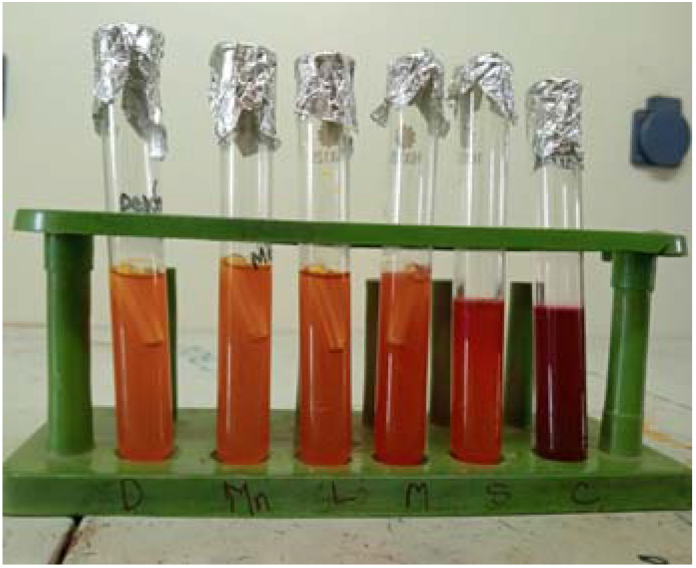
Sugar-fermentation reactions of E. coli isolates with five basic sugars (D = Dextrose, Mn = Mannitol, L = Lactose, M = Maltose, S = Sucrose, C = Control)

**Fig. 10.**
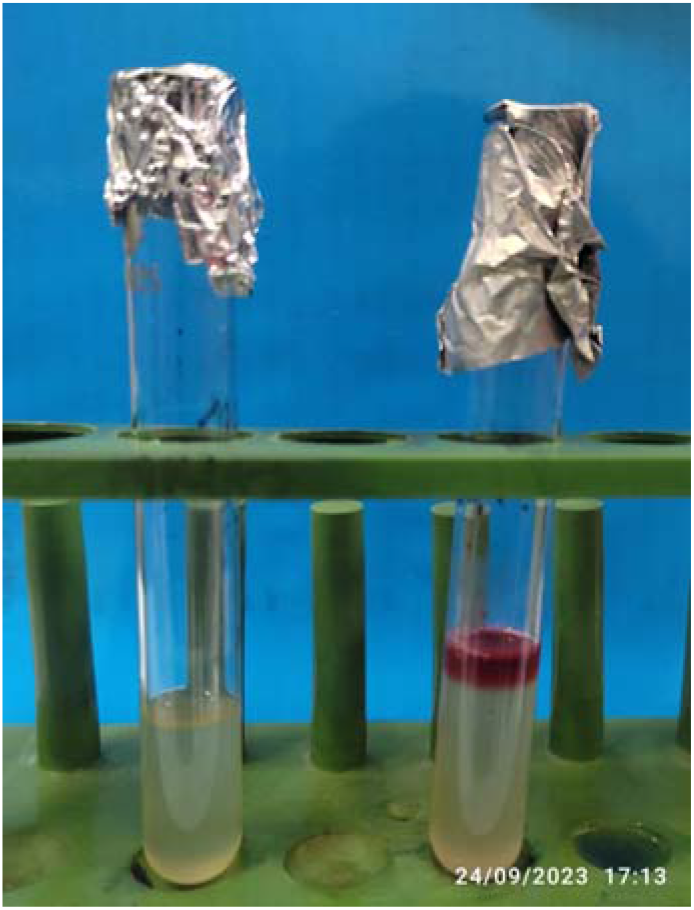
Indole test showing a positive (red ring) and a negative reaction

**Fig. 11.**
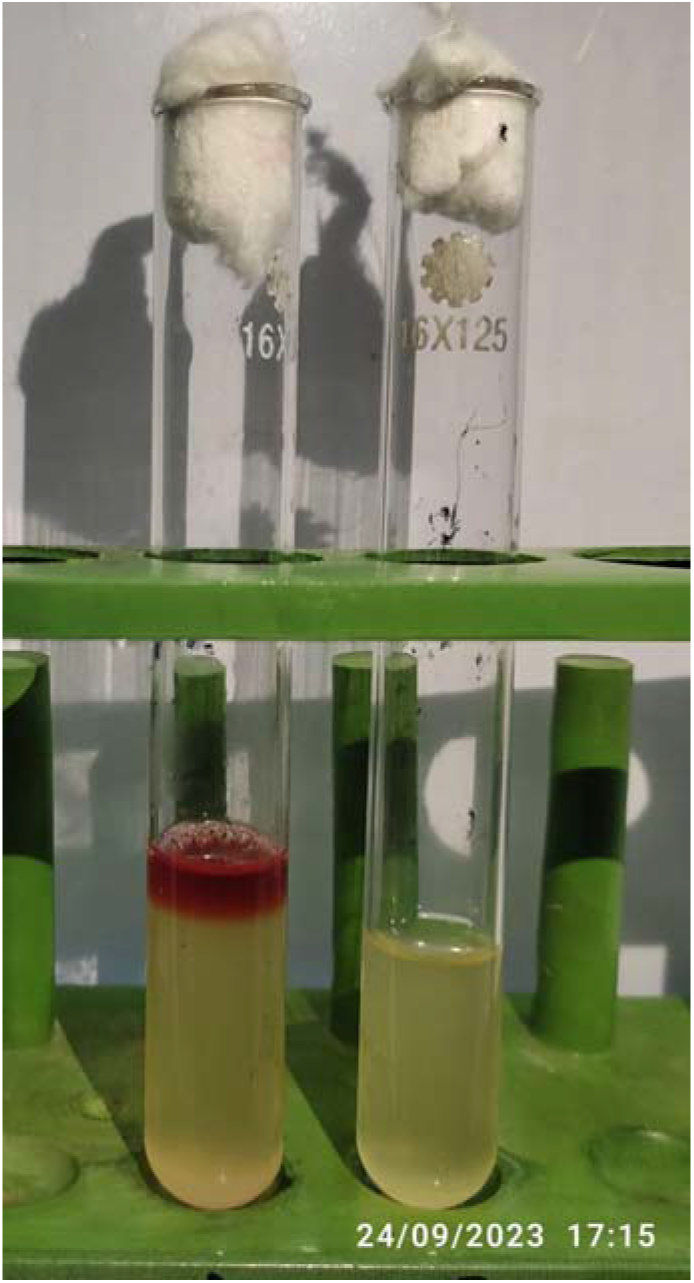
Methyl Red (MR) test showing a positive (red) and a negative (yellow) reaction

**Fig. 12.**
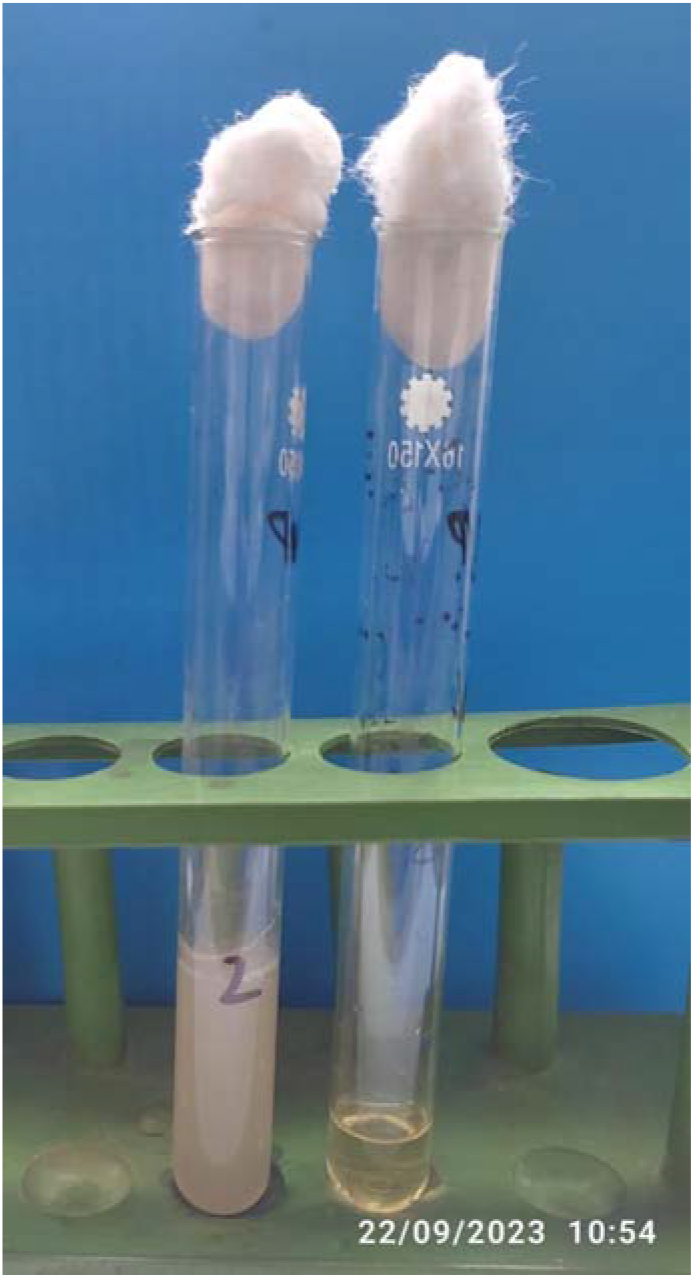
Voges-Proskauer (VP) test showing negative reactions (no pink colouration)

**Fig. 13.**
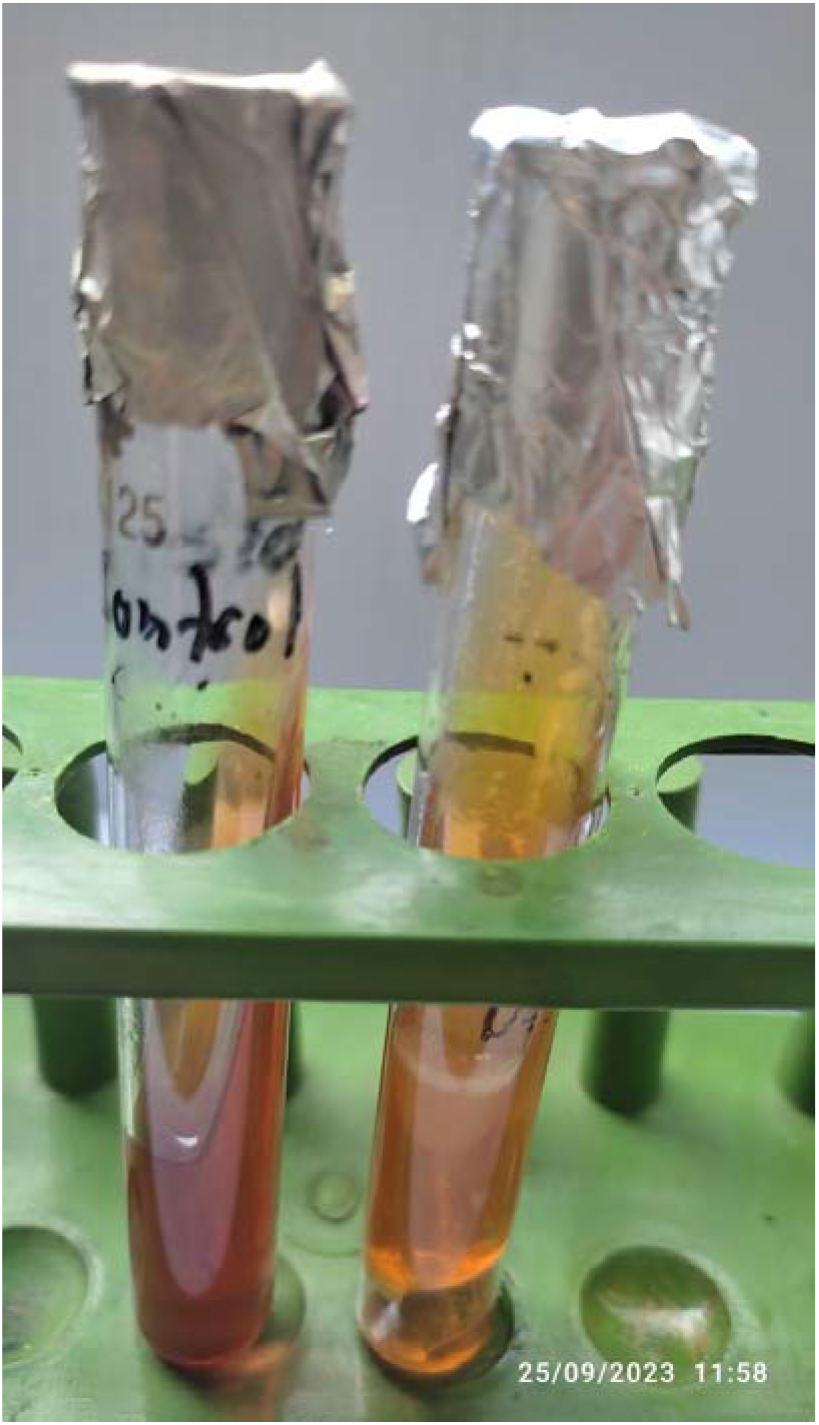
Triple Sugar Iron (TSI) agar slant reaction showing acidic slant and acidic butt with gas production

**Fig. 14.**
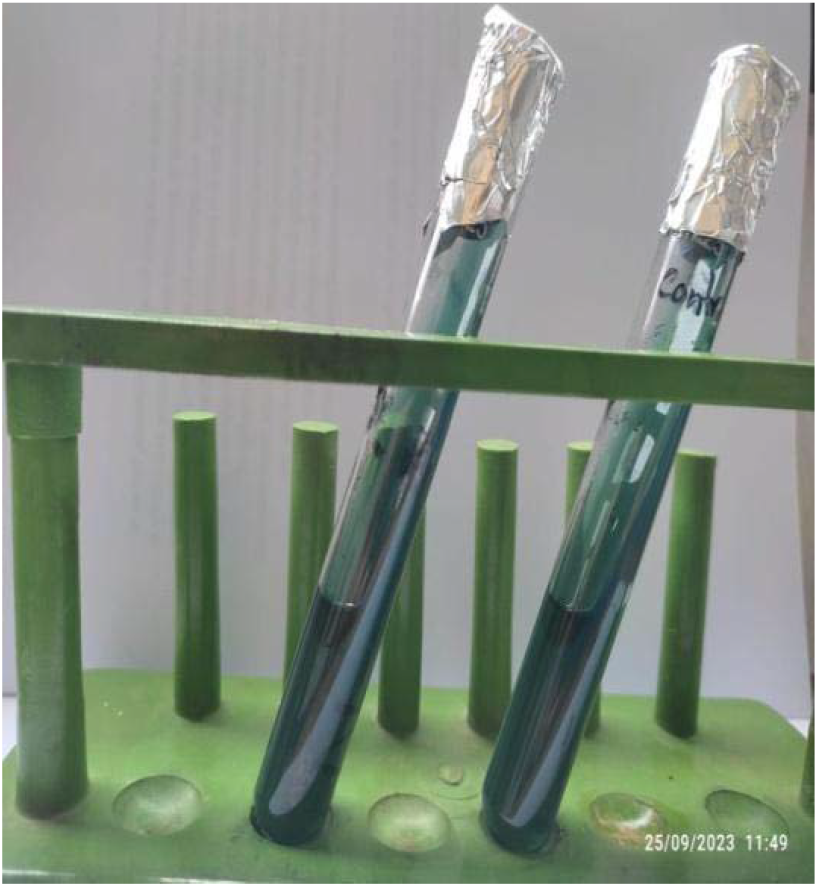
Simmons’ Citrate Utilisation test showing a negative reaction (no colour change)

### 3.4 Prevalence of E. coli in chicken eggs of Natore District

Of the 84 egg samples examined, 56 were positive for *E. coli*, giving an overall prevalence of 66.67%. Prevalence differed among the three sources: 18 of 28 farm-egg samples (64.28%), 16 of 28 market-egg samples (57.14%) and 22 of 28 indigenous-chicken-egg samples (78.57%) were positive (Table 4).

**Table 4.** Prevalence of E. coli in chicken eggs from different sources in Natore District.

| Source of chicken eggs | No. of samples tested | No. positive for *E. coli* | Prevalence (%) | Overall prevalence (%) |
| --- | --- | --- | --- | --- |
| Farm eggs | 28 | 18 | 64.28 |  |
| Market eggs | 28 | 16 | 57.14 |  |
| Indigenous chicken eggs | 28 | 22 | 78.57 |  |
| Total | 84 | 56 | 66.67 | 66.67 |

3.5 **Antibiogram of the isolated E. coli**

Twenty-two confirmed *E. coli* isolates were tested for susceptibility to seven antimicrobial agents by the disc-diffusion method. Resistance was most frequent to neomycin (90.91%) and erythromycin (86.36%), followed by oxytetracycline (77.27%), doxycycline (72.73%), amoxicillin (68.18%), ciprofloxacin (63.63%) and levofloxacin (59.09%). Sensitivity was highest to levofloxacin (27.27%), followed by ciprofloxacin and doxycycline (22.73% each), amoxicillin (18.18%) and oxytetracycline (9.09%); no isolate was sensitive to neomycin or erythromycin. Intermediate sensitivity was most common for doxycycline (40.91%), while the remaining agents showed intermediate sensitivity ranging from 9.09% to 13.63% (Table 5).

**Table 5.** Antibiotic sensitivity and resistance pattern of E. coli isolated from chicken eggs (n = 22)

| Antibiotic | Sensitive n (%) | Intermediate n (%) | Resistant n (%) |
| --- | --- | --- | --- |
| Ciprofloxacin | 5 (22.73) | 3 (13.63) | 14 (63.63) |
| Amoxicillin | 4 (18.18) | 3 (13.63) | 15 (68.18) |
| Doxycycline | 5 (22.73) | 9 (40.91) | 8 (36.36)* |
| Oxytetracycline | 2 (9.09) | 3 (13.63) | 17 (77.27) |
| Levofloxacin | 6 (27.27) | 3 (13.63) | 13 (59.09) |
| Neomycin | 0 (0.00) | 2 (9.09) | 20 (90.91) |
| Erythromycin | 0 (0.00) | 3 (13.63) | 19 (86.36) |

\*As reported in the source data; the sensitive, intermediate and resistant counts for doxycycline are presented as recorded in the original laboratory records.

**Fig. 15.**
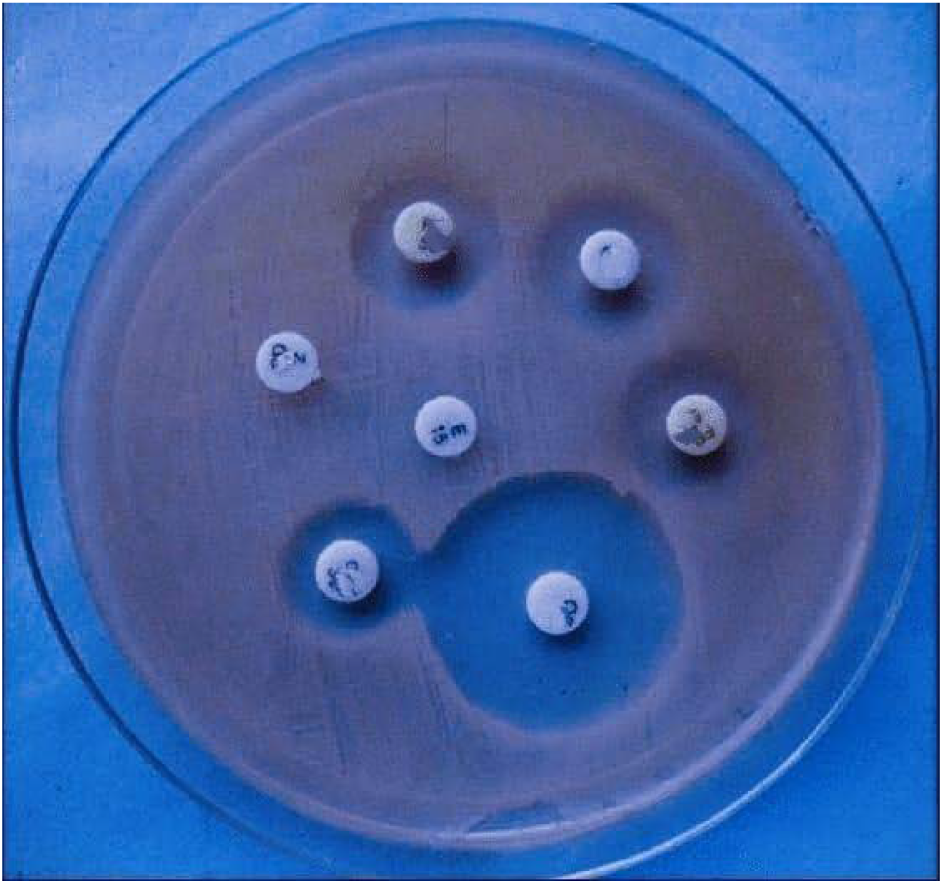
Antibiotic sensitivity (disc-diffusion) test plate showing zones of inhibition around antibiotic discs

## 4. DISCUSSION

Although *E. coli* is a normal commensal of the human and animal intestine, pathogenic strains cause colibacillosis in poultry and vomiting, nausea, diarrhoea and abdominal cramps-occasionally progressing to bowel necrosis-in humans (Todar, 2007). *E. coli* is a Gram-negative, rod-shaped facultative anaerobe (Singleton and Sainsbury, 1981), and in the present study Gram-stained isolates appeared singly or in pairs, consistent with this description.

The overall prevalence of *E. coli* in chicken eggs of Natore District (66.67%) is comparable with the 60.78% prevalence reported in eggs from Australia (Gole et al., 2013) and somewhat higher than the 54.4% prevalence reported in Nigeria (Okorie-Kanu et al., 2016). The 64.28% prevalence recorded in commercial farm eggs in the present study was higher than the 28.7% reported by Abubaker et al. (2018) in similarly sourced eggs, and than the 42% recovery from egg surfaces reported in poultry-farm environments in Bangladesh by Akond et al. (2009). It was also higher than the 21.9% contamination of egg shells reported from Khartoum State, Sudan (Abubaker et al., 2018) and the 25.30% prevalence reported on Ethiopian egg shells (Taddese et al., 2019). Similarly, the prevalence recorded on egg shells in the present study was higher than the 15% reported from Tandojam, Pakistan (Khan et al., 2016), but broadly comparable with the 38.2% contamination of egg shells reported from Enugu State, Nigeria (Okorie-Kanu et al., 2016). These differences between studies most likely reflect variation in hygienic practices during egg handling, storage and marketing, as well as differences in sampling method and season.

The antibiotic-resistance pattern observed in the present study-with resistance to ciprofloxacin, amoxicillin, doxycycline, oxytetracycline, levofloxacin, neomycin and erythromycin at 63.63%, 68.18%, 36.36%, 77.27%, 59.09%, 90.91% and 86.36%, respectively-broadly mirrors the high levels of multidrug resistance reported elsewhere. Adesiyun et al. (2005), using the disc-diffusion method to examine seven treatment agents against 131 *E. coli* and other Enterobacteriaceae isolates, found that 95.4% displayed resistance to one or more antimicrobials, with the highest resistance to streptomycin (90.1%), tetracycline (51.9%) and kanamycin (30.5%). Working with poultry samples from chicken markets in Bangladesh, Akond et al. (2009) reported susceptibility of 86%, 80%, 60%, 36%, 30% and 26% to gentamicin, norfloxacin, ampicillin, tetracycline, neomycin and streptomycin, respectively. Imam et al. (2020) found that 66-100% of *E. coli* strains from Bangladeshi chickens were resistant to tetracycline, penicillin, erythromycin and chloramphenicol, broadly supporting the high resistance to erythromycin and tetracycline-class drugs observed in the present study, whereas Tricia et al. (2006) found no isolate resistant to gentamicin, although 43% were resistant to ampicillin. Such variation in resistance profiles across studies may be explained by differences in the extent of indiscriminate antibiotic use as feed additives, prophylactics or therapeutics, and by differences in prior antimicrobial exposure among the sampled flocks.

Consistent with several previous reports, egg shells were more heavily contaminated with *E. coli* than the internal contents would be expected to be, and the prevalence of *E. coli* on the shells of eggs sampled directly from farms and indigenous households was comparatively high relative to eggs obtained from wholesalers and retail outlets. The degree of risk posed to consumers by contaminated eggs depends on both the type and quantity of bacteria present on the shell and within the egg.

Antibiotic resistance is an increasingly serious problem, particularly in low- and middle-income countries, driven by the overuse of antimicrobials, the intrinsic biological defences of bacteria against antimicrobial agents, and economic and regulatory constraints on rational drug use. Antimicrobial-resistant microorganisms are recognised as one of the foremost threats to global public health, and the present findings-showing very high resistance to neomycin and erythromycin and moderate-to-high resistance to the remaining antibiotics tested-reinforce the need for stronger antimicrobial stewardship in poultry production.

## 5. CONCLUSION

The present study demonstrated a high prevalence of Escherichia coli contamination among chicken eggs collected from commercial farms, markets and indigenous/backyard flocks in Natore District, Bangladesh, with an overall prevalence of 66.67%. The highest prevalence was observed in eggs from indigenous flocks (78.57%), followed by farm eggs (64.28%) and market eggs (57.14%). The isolates were identified based on their characteristic cultural, morphological and biochemical properties. Antimicrobial susceptibility testing revealed substantial resistance to several commonly used antimicrobial agents, particularly neomycin (90.91%), erythromycin (86.36%) and oxytetracycline (77.27%), while resistance was comparatively lower to doxycycline (36.36%) and levofloxacin (59.09%). The high occurrence of E. coli and the observed antimicrobial resistance among egg-associated isolates indicate a potential food-safety and public-health concern. Therefore, improved hygienic practices during egg production, collection, handling, storage and marketing, together with prudent and regulated antimicrobial use in poultry production, are essential. Continued surveillance of E. coli contamination and antimicrobial resistance across the poultry and egg production chain is recommended to support food safety and antimicrobial stewardship.

## Acknowledgements

The authors gratefully acknowledge the Department of Veterinary & Animal Sciences, University of Rajshahi, Bangladesh, for providing laboratory facilities and technical support throughout this study. The authors also sincerely thank the commercial poultry farm owners, egg vendors, and indigenous chicken owners of Natore District for their cooperation during sample collection.

## Ethical Approval

Ethical approval for this study was obtained from the Institutional Animal, Medical Ethics, Biosafety and Biosecurity Committee (IAMEBBC), University of Rajshahi, Bangladesh, before commencement of the study. Egg samples were collected from commercial farms, retail markets, and indigenous chicken flocks with the permission of the respective farm owners, vendors, and household owners. No live animals were experimentally handled or subjected to invasive procedures during this study.

## Informed Consent

Permission was obtained from the respective farm owners, egg vendors, and indigenous chicken owners prior to sample collection. All authors have read and approved the final version of the manuscript and agree to its submission for publication.

## Data Availability

The data supporting the findings of this study are available from the corresponding author upon reasonable request. All data generated or analyzed during this study are included in this article or are available from the corresponding author upon reasonable request.

## Author Contributions

Rubina Khatun: Conceptualization, Literature review, Sample collection, Data curation, Laboratory experiments, Data analysis, Methodology, Statistical analysis, Writing-original draft.

Md. Rimon Bhuiyan: Investigation, Data curation, Data analysis, Laboratory experiments, Literature review, Visualization, Writing-original draft, Writing-review and editing.

Mst. Nahida Akter: Sample collection, Data curation, Laboratory experiments, Investigation, Writing-original draft.

Nise Saha, Sharmin afroz, Anna Purnna Ray: Data curation, Validation, Investigation, Data analysis

K.M. Mozaffor Hossain: Conceptualization, Methodology, Supervision, Project administration, Writing-review and editing, Final approval of the manuscript.

## Funding

This research received no specific grant from any funding agency in the public, commercial, or not-for-profit sectors.

## Conflict of Interest

The authors declare that they have no competing interests or conflict of interest regarding the publication of this manuscript.

## Notes

### Competing Interest Statement

The authors have declared no competing interest.

